# From small to tall: breed-varied household pet dogs can be trained to detect Parkinson’s Disease

**DOI:** 10.1101/2024.01.29.577858

**Authors:** Lisa Holt, Samuel V Johnston

**Affiliations:** PADs for Parkinson’s, 689 Airport Center Road #425, Friday Harbor, WA 98250

**Keywords:** Parkinson’s Disease, medical detection canines, sebum, sniffer dogs, canine olfaction, PADs, Parkinson’s Alert Dogs, Joy Milne

## Abstract

**Objective:** Parkinson’s Disease (PD) is a clinically diagnosed disease that carries a reported misdiagnosis rate of 10–20%. Recent scientific discoveries have provided evidence of volatile organic compounds in sebum that are unique to patients with PD. The purpose of this study was to determine if companion dogs could be trained to distinguish between sebum samples provided by PD-positive patients and PD-negative human controls.

**Methods:** This was a randomized, handler-blind, controlled study. Twenty-three canines of varying breeds, ages, and environmental backgrounds were included. The two-year study period reported here was the final two years of a seven-year program which started in January of 2016. This study encompassed 200 total working session days from 2021 and 2022.

**Results:** When averaged as a group over two years, the 23 dogs were 89% sensitive and 87% specific to an olfactory distinction between PD-positive and PD-negative human donor samples. Ten of the twenty-three dogs averaged 90% or higher in both sensitivity and specificity.

In 161 separate instances, dogs were presented with both unique PD-positive and PD-negative samples (the dogs had not previously encountered any of the samples presented). For these first-time exposures, the 23 dogs collectively averaged 86% sensitivity and 89% specificity.

When comparing the sensitivity and specificity of PD-positive samples from donors who reported levodopa usage to PD-positive samples from donors who reported no levodopa usage, the dogs showed no statistical difference in sensitivity or specificity at the 95% significance level, indicating levodopa usage is not a factor in PD canine detection. Other factors investigated as part of this study included sample donor gender, canine breed, age, duration of time in the program, and training.

**Conclusions:** Companion dogs can be trained with reward-biased detection methodologies to distinguish between PD-positive and PD-negative donor sebum samples in a controlled setting. This study provides further evidence of one or more volatile organic compounds in the sebum of PD-positive patients that can be detected by canines. Summarily, study findings support the application of trained companion dogs for the screening of PD-positive and PD-negative samples in which numbers of samples are limited and the dogs are worked in short intervals, followed by recovery training.

### Box 1. Glossary of Terms

| **Term** | ***As defined and used within the study*** |
| --- | --- |
| **Clear** | An indication of PD-negative (sniffing and not indicating to a sample by the dog) is sometimes referred to as clear or clearing the sample. |
| **Exposure** | A calculation assigned to the number of times a dog exhibits recognition of PD-associated odor and is reinforced for the recognition. |
| **Handler** | The individual responsible for the care and welfare of the dog who also makes the dog available for attendance in working sessions. |
| **Indication** | The manner in which the dog communicates recognition of the target odor, commonly called an “alert.” In this study, dogs were required to hold their indication for a minimum of three seconds in order to receive a positive indication score. |
| **Levodopa** | The most commonly prescribed drug for the treatment of Parkinson’s Disease is levodopa, a central nervous system agent that works to help alleviate some symptoms of PD caused by the reduction of dopamine in the brain. |
| **Negative punishment** | A process of operant conditioning in which the reinforcer (usually food or toy) is withheld from the dog for a behavior in order to decrease the behavior. |
| **Operant conditioning** | In dog training, this refers to the four quadrants of behavior modification that consist of positive reinforcement, negative reinforcement, positive punishment and negative punishment. Dogs in this study were trained with positive reinforcement and negative punishment. |
| **Parkinson’s Disease** | A progressive, incurable disease of the nervous system that causes a deficiency of the neurotransmitter dopamine. |
| **PD-negative** | A sample in which the donor reported no primary associative-PD symptoms and was cleared by 80% of the dogs during the first introduction of the sample. |
| **PD-positive** | A sample in which a minimum of 80% of the dogs had indicated a positive indication during the first introduction of the sample and was also diagnosed by a medical professional. |
| **Positive reinforcement** | A process of operant conditioning in which the dog receives a reinforcer (or reward such as food or toy) for a behavior in order to increase the behavior. |
| **Sensitivity** | The ability to determine a human-sourced sample as positive for PD. |
| **Specificity** | The ability to determine a human-sourced sample as negative for PD. |
| **Target odor** | An odor for which a dog is trained to detect. |
| **Trial** | A presentation of samples to the dogs in which data was collected for the sensitivity and specificity response of each dog to each sample. |
| **Working day** | A day in which the PADs facility was open and available for canine attendance and sample trials. |
| **Working round** | Each working day consisted of two or more rounds. In each round, the dogs were presented with samples in a different order of presentation. |
| **Working session** | The grouped collection of rounds in a working day comprised a working session. |

## INTRODUCTION

Parkinson’s Disease (PD) is a multi-symptomatic progressive neurological disease resulting in fatal symptomatic-system failure. It is estimated that as many as 10 million people worldwide are living with PD (1). In the United States alone, the incidence of PD is estimated at the rate of 90,000 new cases a year. The only neurological disease more prevalent than PD is Alzheimer’s Disease. The greatest known risk factor of PD is aging, with over 90% of cases in the United States at age 65 or older (2). The second-greatest risk factor is gender, with the incidence rate of males developing PD at 1.5× greater than females (2).

There is no conclusive screening test for PD and the diagnosis is most often provided by a combination of clinical assessment of symptoms, dopamine transporter scan, skin biopsy, hereditary, and environmental factors. The misdiagnosis rate in PD is estimated to be between 10–20%, with a greater rate of misdiagnosis occurring within the first two years of diagnosis (3). By the time a patient receives a clinical diagnosis, the disease has progressed to the point of a 60% dopaminergic reduction in the basal ganglia of the brain. Due to this reduction in dopamine, the diagnostic hallmark symptom of tremors become present, along with other associated symptoms, such as loss of balance, cramped handwriting, sleep disturbance, and altered patterns of speech (4).

The gradual reduction of dopamine is estimated to begin in the brain 5–10 years before the presentation of the symptoms that lead to a clinical diagnosis (5). Once this diagnosis has been made, a patient is usually treated with a drug or combination of drugs, which work to cross the blood-brain barrier and deliver a chemical substitute for dopamine, helping to slow, but not stop, the progressive reduction of dopaminergic loss to the brain (6). It is believed that if drugs, along with other therapeutic interventions, could be introduced at the onset of PD, then the progression of the disease could be slowed at a much greater rate than provided through current methods of detection and treatment. This explains the urgent need for an early detectable biomarker of PD (7).

Canines, with their high ability for scent detection, could provide detection for PD-associated odor(s) i.e., a biomarker. However, canines raised in controlled kennel environments for scent detection typically cost between $15,000 and $20,000 per dog. The cost of maintaining a kennel detection environment to raise and train dogs for scent detection is typically over $100,000 per year (8). By contrast, in this study, we set out to explore whether household canine pets could be utilized effectively for the detection of a PD-associated odor, thus eliminating the need for costly, and less available, kennel-raised professional detection dogs.

In 2015, Joy Milne, a woman diagnosed with hyperosmia, a condition that provides her with an extremely sensitive sense of smell, was shown to be able to distinguish between T-shirts worn by PD-positive sample donors and T-shirts worn by human control sample donors (9). The introduction of canine olfaction as a detection tool for the identification of a PD-associated odorant as a biomarker for the disease became a field of canine olfactory study with PADs (Parkinson’s Alert Dogs) for PD located in Washington State. PADs, as an organization, was solely focused on the canine detection of PD and conducted 745 days of research in which data was collected and compiled each day. Our study encompasses the final two years of data from the PADs Program. PADs was the first program to undertake canine detection training for PD-associated odor.

In 2020, working with Joy Milne, researchers at the University of Manchester utilized paper spray ionization mass spectrometry, along with Milne’s sense of smell, to discover more than 500 organic compounds unique to the sebum of PD-positive patients (10). This discovery is being used for the development of a laboratory skin test for the screening of a biomarker for Parkinson’s (9).

Dogs can be trained to detect target odors with a sensitivity that surpasses the capabilities of most modern instruments (11). Dogs can detect odors in parts per trillion (12) and have been successfully trained to use their olfactory ability to correctly identify and distinguish diseases such as lung (13–16), breast (17,18), and colorectal cancers (19), as well as malaria (20), COVID-19 (21–24) and diabetes—each with a reported sensitivity and specificity of >80% (11–14,17,19,20,23,25). In 2022, a study by Gao et al. (26) presented proof of concept for the canine detection of PD in a controlled, laboratory setting. The three canines trained for the study were of the Belgian Malinois breed and were trained in a laboratory setting for canine detection of PD. Before the study, all three dogs had been in training for two years for PD detection. In 2023, a study by Rooney et al (27) provided further supporting evidence of canine detection of PD with findings of two trained dogs in a medical detection program in the U.K.

Household-maintained canines were successfully trained for medical detection purposes in 2008. Medical Detection Dogs, U.K., assigns dogs to handlers who maintain the dog in a home environment, as opposed to a canine group-kenneled environment. The dogs are brought from their home to the facility for training purposes and returned to their home at the end of each training day. The dogs of Medical Detection have been successfully trained for the detection of cancer, diabetes, and malaria. (8)

The importance of breeds for detection has been brought into question in recent years. A canine detection breed comparison study was conducted in which German Shepherds, a breed considered desirable for scent detection, were compared to Pugs, a brachycephalic breed considered unsuitable for scent detection due to the physical attributes of a Pug’s shortened nose. In one study, the Pugs outperformed the German Shepherds in the detection tasks set before them. A third breed, Greyhounds, were also included in the study, but the Greyhounds did not perform well in the scent detection tasks set before them, likely due to lack of interest or drive in the task (28).

Additionally, we wanted to determine if the drug, levodopa, would emerge as a factor in the sensitivity findings of the dogs. Since 1970, levodopa has been the most prescribed drug for the therapeutic treatment of PD (29), and a possibility existed that dogs could target a volatile organic compound as a byproduct of levodopa drug usage, rather than a PD-associated odor(s) caused by the disease.

The primary objective of our study was to determine if detection-trained companion dogs of varied breeds, ages, and backgrounds could distinguish between PD-positive and PD-negative samples. This study further advances the field of canine medical detection of PD by reporting successful findings when the following factors of significance were investigated:

- Levodopa usage as it affects study sensitivity in PD-positive samples.
- Gender differences of sample donors.
- Canine memorization of samples.
- Large variety of breeds, backgrounds and ages of dogs.
- High number of dogs in the trials with no outliers removed.
- High number of trials with no outliers removed.
- Comparison of canine sensitivity between PD-sourced samples that have never been encountered by the dogs to samples that have been previously encountered.
- Length of detection training over extended time.

## METHODS

### Study design

This was a handler-blind, randomized study that included a significant number of detection-trained companion dogs of varying breeds, exposures, and training days to evaluate sensitivity and specificity when presented with PD-positive and PD-negative samples. The study took place in a controlled laboratory setting that was located within a dedicated facility. The facility building was leased and remodeled for the purpose of the seven-year study and included a 14 x 20 ft training area, canine waiting room, research and scribe observation area, sample storage room, office and storage room, and canine exit hallway. The facility was used exclusively for the purpose of this study.

The data collected and reported in this study was compiled from the final two years of a seven-year overall nonprofit canine detection program for the detection of PD. The study period included 200 total working days—95 days in 2021 and 105 days in 2022. Twenty-three dogs participated in a total of 4553 individual trials. The average canine attendance was 8.7 dogs per daily working session. For each daily working session in which 10 or more dogs participated, a minimum of 320 data points were recorded.

Due to a 10%, or higher, potential rate of misdiagnoses in PD, the dogs were at times presented with samples from donors with questionable diagnoses. In these instances, the dogs were not reinforced for a positive indication, regardless of whether it may have been a correct response. This was to reduce the opportunity for any dog to be reinforced for an incorrect response. These samples were only categorized as PD-positive when a minimum of 8 of 10 trained dogs provided a positive indication to the sample when the sample was first introduced. Since no conclusion could be drawn before the analysis of the response of the 10 dogs, no dogs were reinforced until the outcome was determined. This form of withholding the reward carried the risk of extinction of desired response. To counteract this, the dogs underwent training recovery rounds and recovery days on previously determined PD-positive samples, so that the dogs could regain drive for continued work and maintain their expected performance levels for sensitivity and specificity.

If the assessment sample in the first round of presentation to the dogs was deemed PD-positive based on the response of 80% or higher sensitivity of 10 dogs, then the assessment sample was then presented in a second round, in a new position on the wheel (see Figure 2), and the dogs were reinforced for a correct response in that round. If the assessment sample in round one was deemed PD-negative based on the response of 79% or lower sensitivity of 10 dogs, then the dogs were presented with a different, previously assessed PD-positive sample, in the second round.

**Figure 1.**
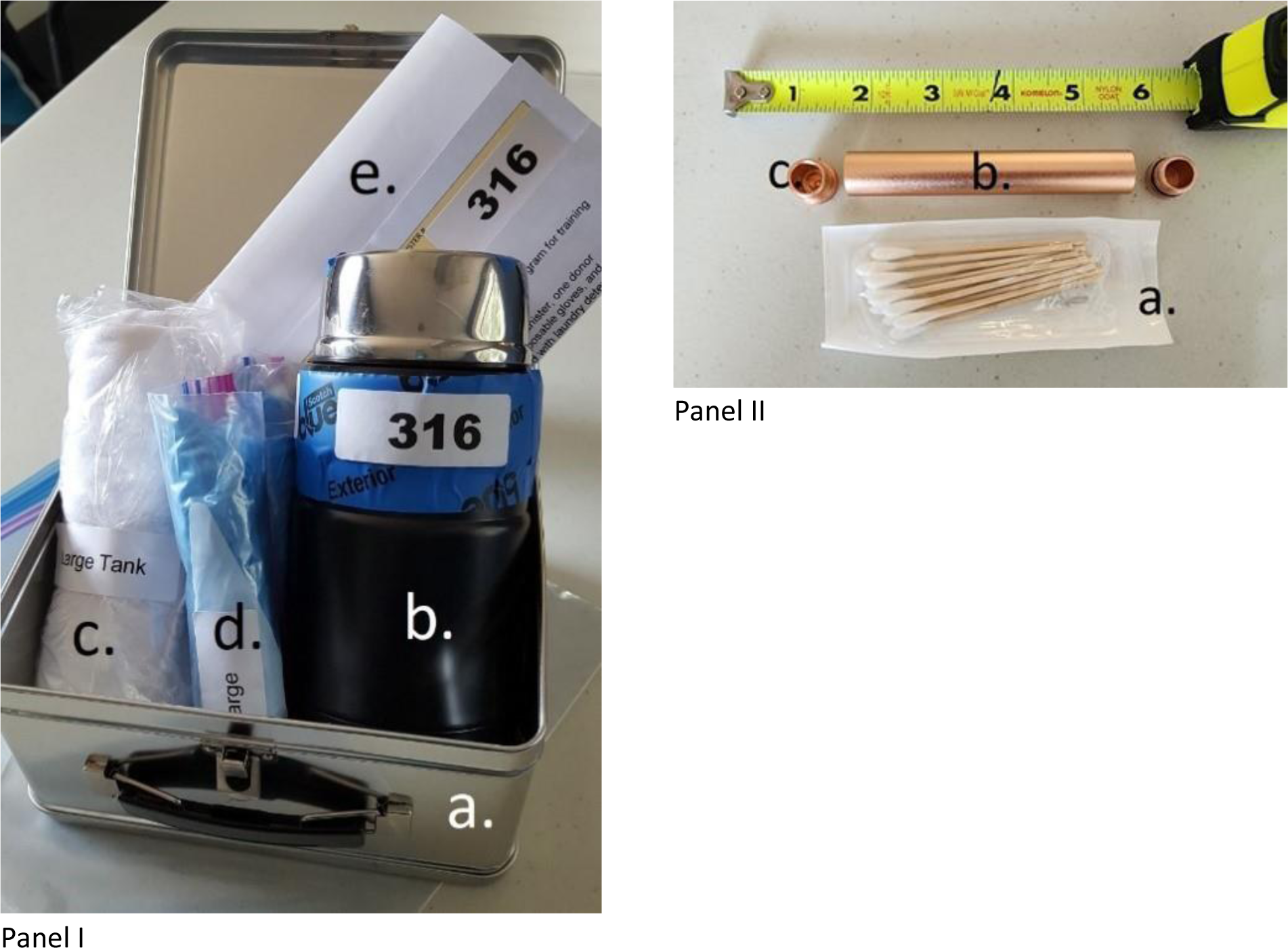
Sample kit including T-shirt and canister, with additional materials for swab sample collection. Panel I: a) Kit container, b) Sample container, c) T-shirt sample to be worn by donor, d) Disposable gloves for caretaker (if assistance is needed), e) Research collection forms; Panel II: a) Sterile swabs, b) Swab sample container, c) Removable cap at either end so that heads of swabs are available for sniffing without being handled by a researcher.

**Figure 2.**
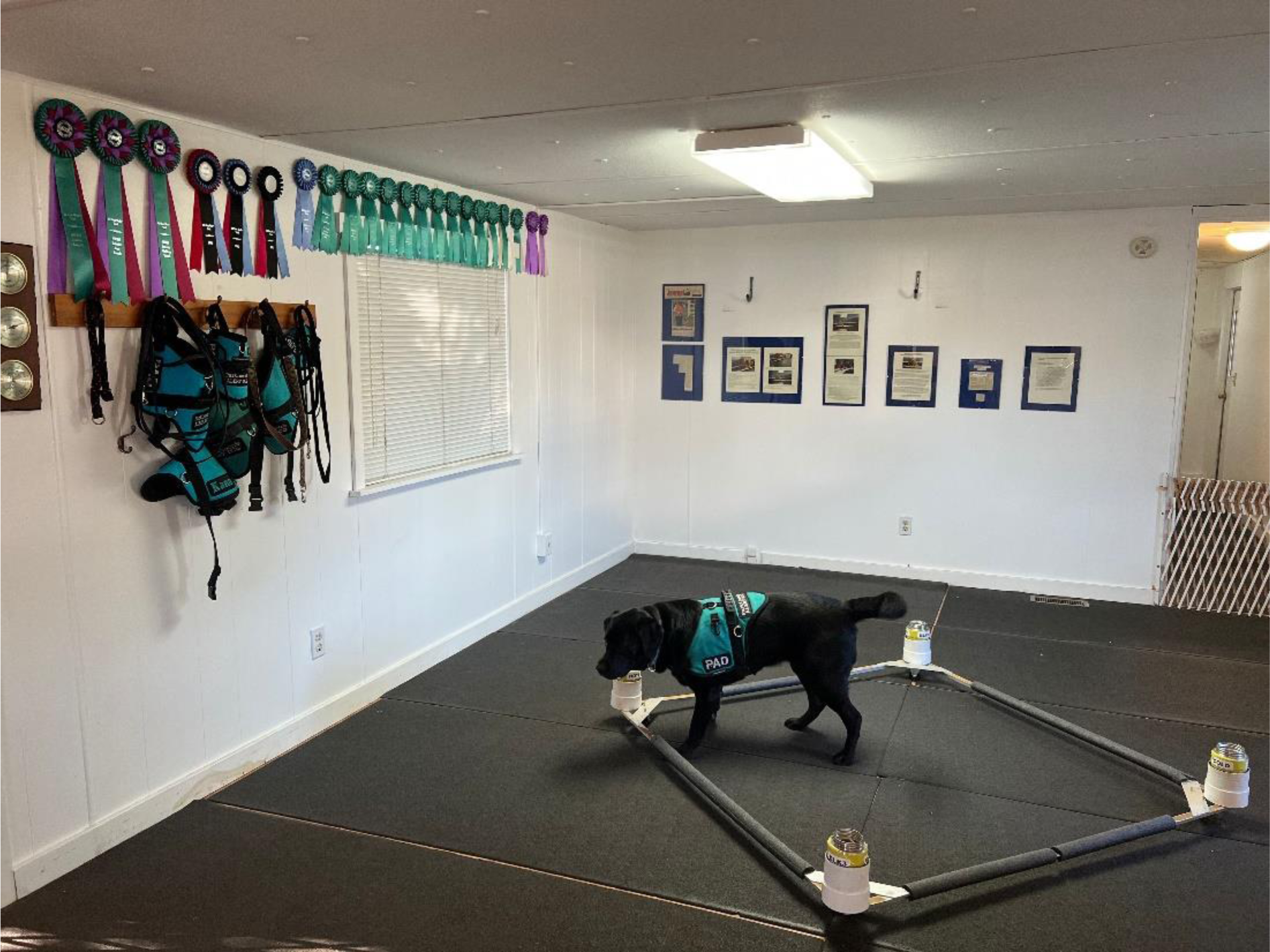
Sample presentation wheel. Sample canister holders were elevated above the flooring to prevent residual odor contamination to the floor. The wheel is designed to be rotated in any direction.

In instances where the dogs were not reinforced for correct sensitivity responses on PD-positive samples (i.e. assessments), the data were not included in the final statistical analysis, but specificity on the PD-negative samples was included. This was because, in assessment sessions with the assessment sample of unknown PD status, the human control sample was a known PD-negative sample.

### Sample participants

PD-positive sample donors were recruited from Parkinson’s support groups, Rock Steady Boxing (a PD-participant fitness organization), physicians, and neurologists, as well as via media coverage and pull-through responses from the website, padsforparkinsons.org). Control sample donors were recruited from activity clubs, friends, and family members of program volunteers, businesses, and other events. All samples were collected from donors who provided informed consent, and personal donor sample information was blinded from all study personnel, affiliates, and media. The informed consent document used to obtain permission for sample inclusion in this research study was prepared and reviewed by legal counsel.

There were 43 total PD-positive sample donors and 31 total PD-negative sample donors whose samples were used in this study spanning 2021 and 2022 (Table 1). PD-participant donors were screened for levodopa drug usage, age, gender, onset of symptoms, type of diagnosis, date of diagnosis, and whether the diagnosis was by a physician, movement disorder specialist, or neurologist. Overall, 24 of 43 sample donors were diagnosed by a neurologist, one was diagnosed by a movement disorder specialist, and the remaining 18 sample donors were diagnosed by a physician. Control participants were screened for age and gender and were eliminated for any of the following: household partner diagnosed with PD, anosmia, bowel disorder, sleep disorder, changes in gait, speech, or handwriting. When all unique samples were presented to the dogs, the Control and PD samples were matched in age within 10 years and matched by gender in 80% or greater instances. Both PD participant and Control participant donor samples were of varied geographic origin within the United States. Table 2 lists the control sample donor characteristics for the 31 control samples used in this study.

**Table 1.** PD-positive sample donor characteristics by age group and levodopa usage.

| Age Bin | Sample Size | % Male | % Female | % levodopa Use |
| --- | --- | --- | --- | --- |
| <= 50 | 1 | 0 | 100 | 0 |
| 51–60 | 5 | 60 | 40 | 80 |
| 61–70 | 14 | 71 | 29 | 57 |
| 71–80 | 20 | 70 | 30 | 55* |
| 81–90 | 3 | 33 | 67 | 67 |
| Total | 43 | 65% | 35% | 58%* |

\*One sample donor indicated levodopa usage was unknown

**Table 2.** PD-negative donor sample characteristics by age group.

| Age Bin | Sample Size | % Male | % Female |
| --- | --- | --- | --- |
| <=30 | 12 | 50 | 50 |
| 31–40 | 6 | 50 | 50 |
| 41–50 | 5 | 60 | 40 |
| 51–60 | 5 | 60 | 40 |
| 61–70 | 3 | 67 | 33 |
| Total | 31 | 55% | 45% |

For this two-year study, 58% of PD sample donors were selected based on confirmed diagnosis by a neurologist and the elapsed time since diagnosis and symptoms. The remaining 42% of the PD-patient sample donors were selected based on reported symptoms with clinical diagnoses. The levodopa-negative Parkinson’s sample donors comprised 17 of the 43 total sample donors. Control sample donors were of varying age, ethnicity, and gender, reporting no associative Parkinson’s symptoms.

### Sample material

T-shirts that were worn overnight by all participants were chosen as sample material. This selection was based on the Joy Milne test developed by Dr. Tilo Kunath at the University of Edinburgh (9). All T-shirts provided to both PD and control participants were purchased from the same manufacturer, tank style, ribbed, 100% cotton, and washed under the same conditions in the same machine, packaged and shipped using the same materials. Along with a new, freshly washed T-shirt, sample donors were provided with a new, stainless-steel double-walled vacuum-insulated 24-ounce thermos and a new metal lunchpail for packaging the T-shirt, once worn, for the return to the program (Figure 1). The thermos was designed to hold heat for 14 hours and cold for up to 24 hours and was selected as a solution for containing volatile organic compounds within a container for shipping and storage. Donor sample participants were instructed to pack the T-shirt into the thermos with the neck of the T-shirt at the top of the thermos.

### Sample storage

All thermoses containing samples were stored in a dedicated cooler set at 11 °C and presented to the dogs within 30 days of receipt. Samples on which the dogs performed with averages of 80% or higher sensitivity as indicated by 10 or more dogs in one session were used for maintenance training, recovery training, and foundation training of new dogs.

In 2022, we learned that Medical Detection Dogs, U.K., had successfully trained dogs for the detection of PD-associated odor using cotton swabs. Based on this information, in September of 2022, the PADs program changed the sample material used in canine detection from T-shirts to cotton swabs, a less expensive and more efficient sample material for collection and storage. Under the sample collection protocol for swabs, 10 cotton swab samples were collected from the upper back and neck area of each PD and control sample participant. The swab samples were collected by a gloved PADs researcher and each sample collection of 10 swabs was placed in a 4.3” height × ½” diameter metal tube that was capped at both ends. This allowed for the swab-tip end of the cotton swab to be opened for sniffing by the dogs without handling by a researcher. The tubes, containing swab samples, were stored under the same protocol, and handled under the same procedure as the T-shirt samples (Figure 1).

### Canine participants

A total of 23 canines of various breeds and ages participated. The canines were selected based on age, demonstration of drive, and availability for training, with little consideration given to breed or sex. All dogs completed a minimum of eight months of prior training for PD detection, whereas some had completed as much as five years of prior training. The Parkinson’s canine detection program had been underway since 2016, so all dogs in the study were previously trained to distinguish between T-shirt samples worn by Parkinson’s patients and T-shirt samples worn by healthy human controls with an average of 75% or higher sensitivity and specificity. Breeds represented in the study included commonly utilized breeds for detection purposes, such as Labrador Retriever and Vizsla, and some of the dogs were of less commonly utilized breeds, such as Pomeranian and English Mastiff. Table 3 shows the breed, age, and sex of each dog at the beginning of the study.

**Table 3.**
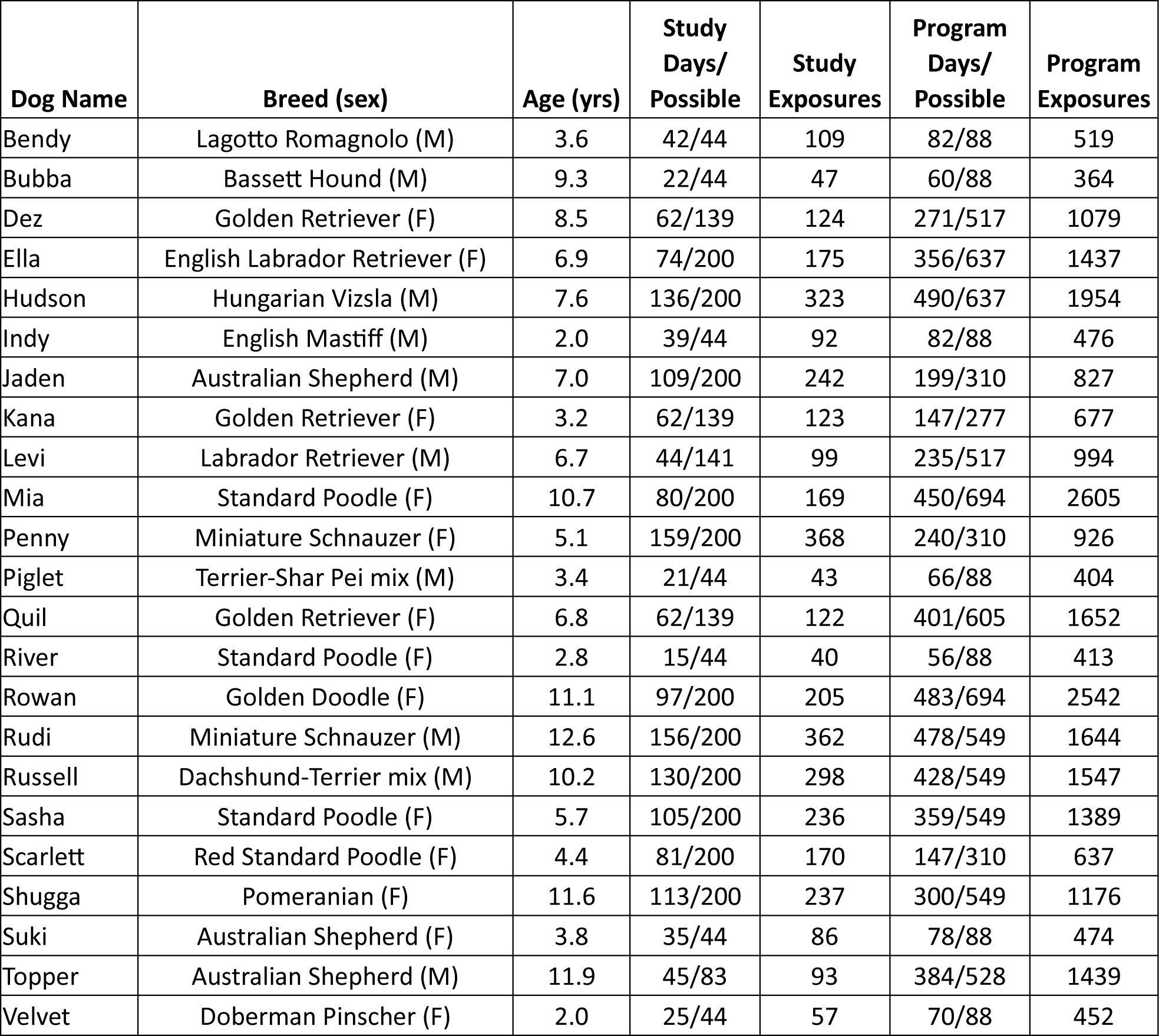
Characteristics of all canine participants in this study.

| Dog Name | Breed (sex) | Age (yrs) | Study Days/<br>Possible | Study Exposures | Program Days/<br>Possible | Program Exposures |
| --- | --- | --- | --- | --- | --- | --- |
| Bendy | Lagotto Romagnolo (M) | 3.6 | 42/44 | 109 | 82/88 | 519 |
| Bubba | Basset Hound (M) | 9.3 | 22/44 | 47 | 60/88 | 364 |
| Dez | Golden Retriever (F) | 8.5 | 62/139 | 124 | 271/517 | 1079 |
| Ella | English Labrador Retriever (F) | 6.9 | 74/200 | 175 | 356/637 | 1437 |
| Hudson | Hungarian Vizsla (M) | 7.6 | 136/200 | 323 | 490/637 | 1954 |
| Indy | English Mastiff (M) | 2.0 | 39/44 | 92 | 82/88 | 476 |
| Jaden | Australian Shepherd (M) | 7.0 | 109/200 | 242 | 199/310 | 827 |
| Kana | Golden Retriever (F) | 3.2 | 62/139 | 123 | 147/277 | 677 |
| Levi | Labrador Retriever (M) | 6.7 | 44/141 | 99 | 235/517 | 994 |
| Mia | Standard Poodle (F) | 10.7 | 80/200 | 169 | 450/694 | 2605 |
| Penny | Miniature Schnauzer (F) | 5.1 | 159/200 | 368 | 240/310 | 926 |
| Piglet | Terrier-Shar Pei mix (M) | 3.4 | 21/44 | 43 | 66/88 | 404 |
| Quil | Golden Retriever (F) | 6.8 | 62/139 | 122 | 401/605 | 1652 |
| River | Standard Poodle (F) | 2.8 | 15/44 | 40 | 56/88 | 413 |
| Rowan | Golden Doodle (F) | 11.1 | 97/200 | 205 | 483/694 | 2542 |
| Rudi | Miniature Schnauzer (M) | 12.6 | 156/200 | 362 | 478/549 | 1644 |
| Russell | Dachshund-Terrier mix (M) | 10.2 | 130/200 | 298 | 428/549 | 1547 |
| Sasha | Standard Poodle (F) | 5.7 | 105/200 | 236 | 359/549 | 1389 |
| Scarlett | Red Standard Poodle (F) | 4.4 | 81/200 | 170 | 147/310 | 637 |
| Shugga | Pomeranian (F) | 11.6 | 113/200 | 237 | 300/549 | 1176 |
| Suki | Australian Shepherd (F) | 3.8 | 35/44 | 86 | 78/88 | 474 |
| Topper | Australian Shepherd (M) | 11.9 | 45/83 | 93 | 384/528 | 1439 |
| Velvet | Doberman Pinscher (F) | 2.0 | 25/44 | 57 | 70/88 | 452 |

### Foundation detection training

Before study selection and during the study period, all dogs were exposed to operant conditioning training methods in their home environment. Owner/handlers engaged in primarily two quadrants of behavioral training by utilizing positive reinforcement and negative-punishment methods for shaping canine behavior.

All dogs were trained for PD odor detection by a single detection trainer. None of the dogs were owned or separately handled by the detection trainer. The dogs underwent foundation training through methods similar to controlled substance detection training, beginning with building hunt drive on a primary reinforcer, then followed by pairing in which the primary reinforcer was available to the dog along with the target odor. At the final stage of foundation training, the dogs worked in the absence of pairing and received a reinforcer for an indication of the target odor. During foundation training, all dogs underwent a minimum of 300 paired exposures (the primary reinforcer, i.e., food, was available to the dog at the time of exposure) at the rate of 10 exposures per training day on their target odor for 6 months. The dogs were also subjected to varied background noise in both target and control samples and were presented with distractors (i.e., onion, coffee, spices, soaps, etc.) and clear (absent of target odor) rounds. In clear rounds, where no target odor was presented, the dogs were trained to go to the exit gate of the facility and were then reinforced outside of the exit gate. During foundation training, all dogs were presented with equal numbers of human-sourced control samples and PD-positive samples so the dogs would learn to distinguish their target odor from the background noise of human scent.

Once a dog had detected the target odor, the dog was trained to hold its body position at the target odor for a minimum of three seconds before receiving a reinforcer. This allowed adequate time for the dogs to clear the control samples with most clearing of samples taking place in one second or less. The dogs were allowed to sit, stand, down, or point while holding, but an alert was not confirmed unless the dog held the position for three or more seconds (30). If a dog held position at a human-sourced control sample for three seconds and then moved onto the target odor for three seconds, the dog was reinforced for indication of the target odor but scored as 0% sensitive, 0% specific for the round (see Table 4, Scoring of the Dogs).

**Table 4.**
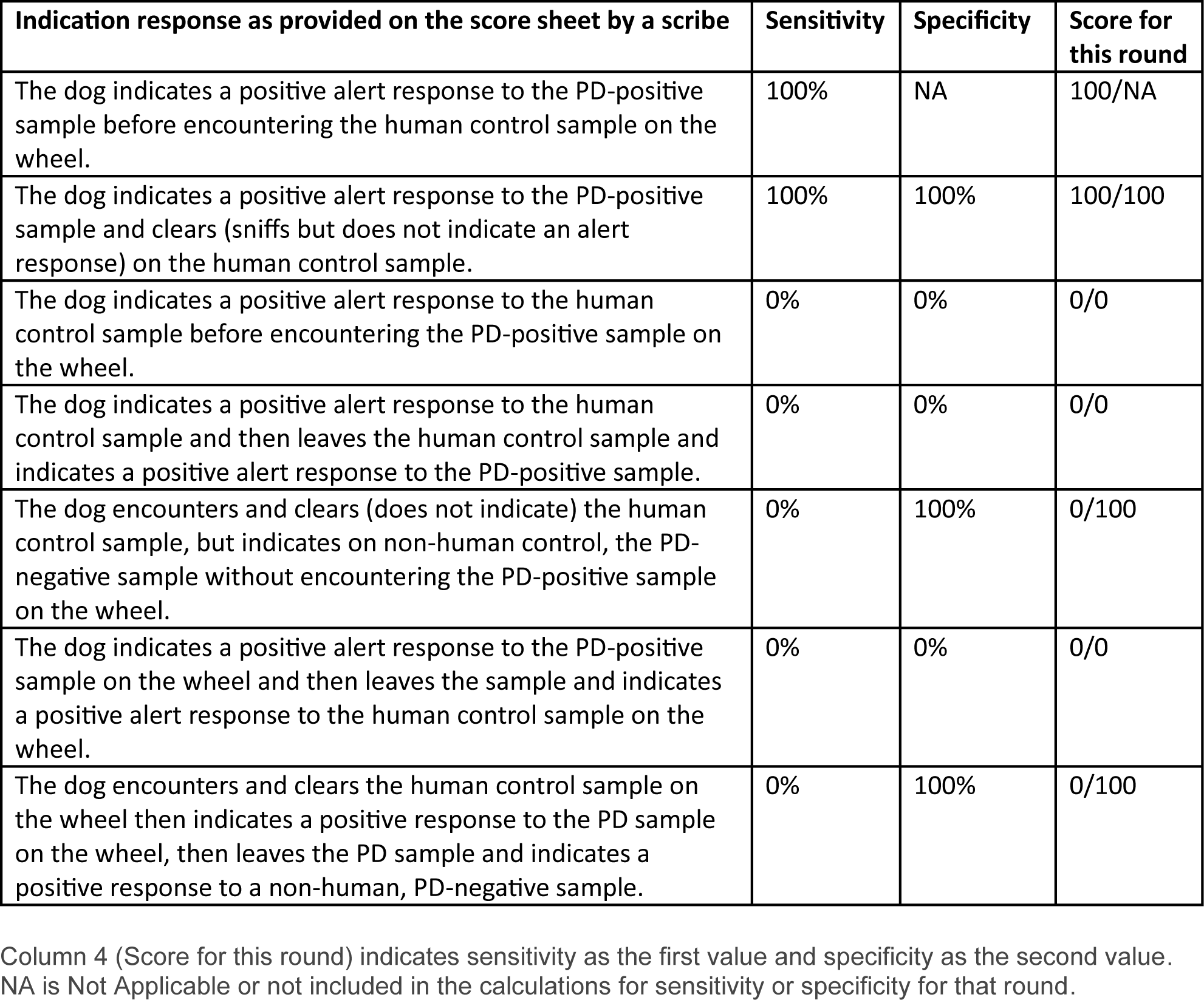
Scoring of the dogs for each round.

Once foundation training had been completed and a dog was able to demonstrate the olfactory distinction between a unique unpaired sample containing the PD-associated target odor (sensitivity) and that of a human control sample (specificity) of 80% or higher across six working sessions on varied samples, the dog was considered trained to detect PD-associated odor and suitable for study inclusion.

### Sample presentation

The donor T-shirt or swab samples within the stainless-steel canisters were not handled by the researcher or any member of the research team. A gloved researcher placed each sample canister on the wheel as indicated by color-coded random selection. The selector indicated the position of each sample to be placed on the wheel, and whether the wheel was to be rotated in counterclockwise or clockwise position between rounds. The wheel was rotated into the position as indicated by the randomizer, and then the predetermined PD-negative control samples were taken from the refrigerated storage unit, canister lids were uncapped, and the control samples were placed onto the wheel (Figure 2). The trainer re-gloved and then placed the PD-positive sample last on the wheel. The canister holders on the wheel were routinely wiped with isopropanol to eliminate lingering or residual odor.

For sniffing indication by the dogs, four samples were randomly placed on a sample wheel. To minimize the potential for fringing (an alert that occurs when the dog indicates close to the target odor but not at the source) and reduce odor transfer between samples, the canisters were spaced four feet (1.23 m). The wheel, for each round of canine sniffing, held one PD-positive sample, one human-sourced control sample, and two other unworn T-shirt samples. The wheel position in the room, sample position on the wheel, and order of dogs were all placed by random selection. The dogs worked off-leash and were free roaming as they entered the sample presentation room and were permitted to work the wheel in any direction. Some of the dogs worked the wheel in a clockwise direction and some worked counterclockwise. A few of the dogs were prone to changing direction based on the scent signature in the room. The dogs worked individually, one after the other, for two or three consecutive rounds and maintained the same run order as determined for the day. The sample wheel was rotated for each round so that, for each round, the dogs encountered the samples in a different position on the wheel.

For most of the study period, there were four working days per week. Not all dogs attended each working day, but most dogs attended at least one or two working days per week. This means that, on each working day, there was a different assembly of dogs. The dogs also varied in their run order, so that the order in which the dogs encountered the samples on the wheel and were subsequently scored was random on each day attended. The dogs were further varied in their ages, breeds, environmental and genetic factors, sex, duration, and experience within the program.

### Handler-blind research protocol

All handlers observed the dogs under blind conditions and were instructed to stand motionless in one position outside of the immediate training area. One trainer/researcher observed the dogs in a mirror while standing motionless in the same marked spot through all rounds and all sessions in the study (Figure 2). While the dogs sniffed samples, no eye contact was made by any researcher or trainer with the dogs. A data recorder (scribe) sat outside the training room behind a half-wall barrier and was able to observe the dogs from a distance.

### Scoring of the dogs

Indications were confirmed between the trainer and scribe and determined as 100% sensitive if the dog’s first 3-second indication (hold) on a PD participant-sourced sample was considered correct, and the dog had not held a three-second alert on any other sample on the wheel. If the dog cleared the human-sourced PD-negative control sample (sniffed and did not hold position) on the wheel before indicating on the PD-positive sample, the dog was determined as 100% specific for the round. If the dog indicated the target sample as positive before encountering the human control sample on the wheel, the specificity for that round was scored as “NA” or non-applicable and did not apply to the daily specificity calculations for the dog or sample in that round. If a dog were to hold for three seconds on a human-sourced control sample at any time in the round, the dog was scored as 0% specific and 0% sensitive. In this instance, the dog was scored as 0% sensitive, even if the dog did not have the opportunity to sniff the PD-sourced sample. (See Table 4 for an explanation of scoring).

Sensitivity and specificity were both calculated as in Trevethan (31). The overall averaged specificity with all control samples included (human-sourced and non-human-sourced controls) was 91.2%. When the non-human sourced controls were eliminated from the specificity calculations, specificity was at 86.6%, a truer calculation of PD-positive human-sourced samples as compared to PD-negative human-sourced samples as per the objective of the study. In all instances, a human-sourced control sample was presented on the wheel. In this manner, the dogs were scored for sensitivity and specificity to mimic a correct or incorrect response in a simulated screening scenario in which all people, or all samples, would exude human scent signature.

### Statistical methods

Sensitivity and specificity data were recorded for each dog on prepared data sheets as the dog exited the training area. These daily rounds data sheets were then manually summarized into daily summaries, and then into monthly canine performance reports. The data from the daily sheets were also entered into individual daily spreadsheets. Daily spreadsheets were then combined and summarized to match the manually summarized reports, then verified against the manually entered canine performance reports, and discrepancies were corrected to match the original recordings made during each working session.

The verified spreadsheet data sets were imported into Microsoft Access database tables, where queries were developed to select and summarize the results. Where confidence limits are presented, the Clopper-Pearson (exact) confidence interval method (32) for binomial trials was used through the confidence interval for a proportion (33). Where comparisons of proportions are presented, the N-1 Chi-squared test as recommended by Campbell (34) and Richardson (35) was employed in MedCalc Software Ltd. For the comparison of proportion calculators (36) of levodopa usage results, a two-proportion z-test was used (37).

## RESULTS

### Average sensitivity/specificity

Table 5 summarizes the averaged sensitivity and specificity by dog for the two years of the study (4959 total dog-sample encounters). Assessment days were included if the sample was found to be PD-positive. During assessment days, sensitivity was not recorded, but specificity was. The dogs are ranked by percent sensitivity, excluding assessment days. There were 10 “top-tier” dogs (90% or higher in both sensitivity and specificity), although three of those dogs had less than 100 rounds.

**Table 5.** Averaged sensitivity and specificity for 23 dogs for January 2021—November 15, 2022 (top-tier dogs in bold).

| Dog Name | % Sensitivity | Indications (Correct/Total) | % Specificity | Indications (Correct/Total) | Attendance Days/ Days Possible |
| --- | --- | --- | --- | --- | --- |
| **Scarlett** | 97.55 | (199/204) | 97.10 | (134/138) | (81/200) |
| **River** | 95.45 | (42/44) | 95.65 | (22/23) | (15/44) |
| **Russell** | 94.63 | (335/354) | 94.15 | (193/205) | (130/200) |
| **Quil** | 93.63 | (147/157) | 94.03 | (126/134) | (62/139) |
| **Hudson** | 93.57 | (349/373) | 90.43 | (208/230) | (136/200) |
| **Jaden** | 92.91 | (262/282) | 94.44 | (153/162) | (109/200) |
| **Penny** | 92.61 | (401/433) | 93.01 | (253/272) | (159/200) |
| **Velvet** | 92.54 | (62/67) | 92.86 | (52/56) | (25/44) |
| **Bendy** | 91.74 | (111/121) | 90.24 | (74/82) | (42/44) |
| **Sasha** | 91.73 | (255/278) | 93.31 | (237/254) | (105/200) |
| Ella | 91.19 | (176/193) | 87.10 | (108/124) | (74/200) |
| Suki | 89.80 | (88/98) | 91.89 | (68/74) | (35/44) |
| Kana | 89.17 | (140/157) | 87.93 | (102/116) | (62/139) |
| Rowan | 88.80 | (222/250) | 87.30 | (165/189) | (97/200) |
| Topper | 88.70 | (102/115) | 80.33 | (49/61) | (45/83) |
| Levi | 88.03 | (103/117) | 83.33 | (65/78) | (44/141) |
| Mia | 85.85 | (176/205) | 79.20 | (99/125) | (80/200) |
| Dez | 82.17 | (129/157) | 71.26 | (62/87) | (62/139) |
| Rudi | 81.97 | (350/427) | 75.10 | (193/257) | (156/200) |
| Shugga | 81.54 | (243/298) | 73.30 | (129/176) | (113/200) |
| Indy | 76.64 | (82/107) | 72.06 | (49/68) | (39/44) |
| Piglet | 74.55 | (41/55) | 71.43 | (25/35) | (21/44) |
| Bubba | 62.30 | (38/61) | 55.32 | (26/47) | (22/44) |
| **Average** | **89.02%** | **(4053/4553)** | **86.60%** | **(2592/2993)** |  |

For all 23 dogs in the program, the combined overall average sensitivity was 89.0% (4053 correct / 4553 total encounters), and overall specificity was 86.6% (2592 passed-up human control / 2993 total human control encounters). Specificity includes a lower number of encounters because sometimes dogs would alert on the PD sample before encountering the human control sample.

### Average sensitivity/specificity for top-tier dogs

For dogs that achieved 90% or higher for both sensitivity and specificity (10 dogs: Sasha, Bendy, Velvet, Penny, Jaden, Hudson, Quil, Russell, River, Scarlett), the overall sensitivity was 93.5% (2163 correct / 2313 total encounters) and the overall specificity was 93.3% (1452 passed up human control / 1556 total human control encounters).

### Average sensitivity and specificity when encountering all unique PD samples and control donor samples

The average sensitivity for first-time encounters (first round only with no warm-up round, so all first encounters were cold runs) with a unique PD sample and unique human control sample was 86.3% (139 correct / 161 total encounters). The average specificity for first-time encounters (first round only) with a unique PD sample and unique human control sample was 89.0% (121 passed-up human control / 136 total human control encounters). Dogs encountered the unique samples in random order for each round, and the samples were in randomized positions in the room for each round.

Comparing the sensitivity and specificity for first-time encounters with a unique PD sample and unique human control sample to the overall sensitivity and specificity for first-round encounters (with unique sample rounds removed) showed no significant difference (p = 0.95) as shown in Table 6. Specificity was higher for those rounds with unique samples but did not exceed the 90% confidence level. This indicates that it is not likely that the dogs are indicating PD-positive samples by memory.

**Table 6.**
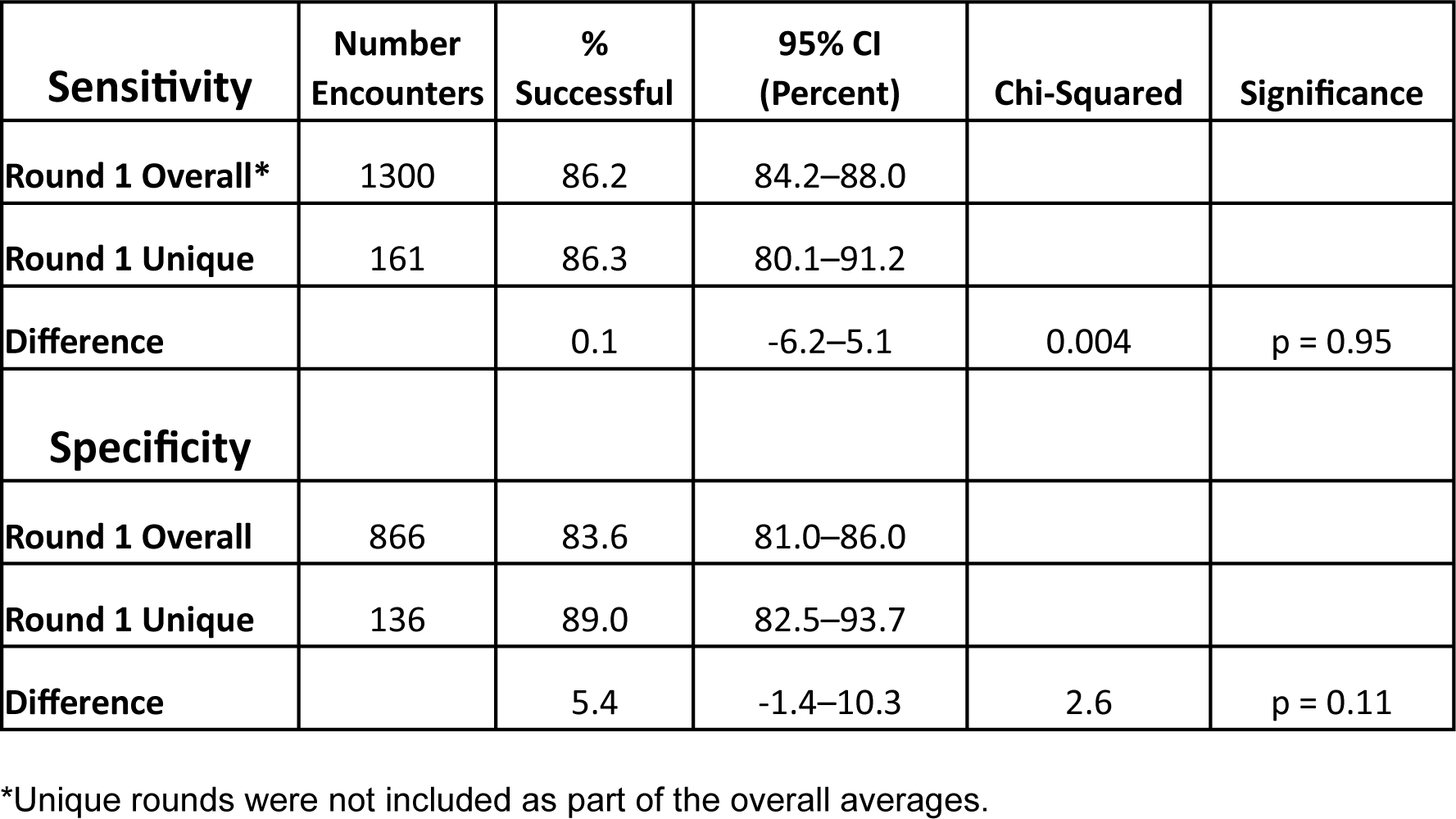
Comparison of Overall Round 1 encounters (unique encounters removed) with Unique Round 1 encounters.

| **Sensitivity** | **Number Encounters** | **% Successful** | **95% CI (Percent)** | **Chi-Squared** | **Significance** |
| --- | --- | --- | --- | --- | --- |
| **Round 1 Overall\*** | 1300 | 86.2 | 84.2–88.0 |  |  |
| **Round 1 Unique** | 161 | 86.3 | 80.1–91.2 |  |  |
| **Difference** | | 0.1 | -6.2–5.1 | 0.004 | $p = 0.95$ |
| **Specificity** |  |  |  |  |  |
| **Round 1 Overall** | 866 | 83.6 | 81.0–86.0 |  |  |
| **Round 1 Unique** | 136 | 89.0 | 82.5–93.7 |  |  |
| **Difference** | | 5.4 | -1.4–10.3 | 2.6 | $p = 0.11$ |

\*Unique rounds were not included as part of the overall averages.

### Average sensitivity/specificity for top-tier dogs when encountering all unique PD and control donor samples

The combined sensitivity for the 10 top-tier dogs (90% overall sensitivity and specificity for the entire study period) when they first encountered a unique PD and unique human control sample (first round only) was 87.1% (74 correct / 85 total encounters). The combined specificity for the same group and samples was 90.4% (66 passed-up human control / 73 total human control encounters).

### Averaged sensitivity/specificity for rounds 1 and 3

The overall combined average sensitivity for all dogs in round 1 of each daily session (where both round 1 and round 3 were attended) was 86.2% (1174 correct / 1362 total encounters), whereas the overall combined average sensitivity for all dogs in round 3 of each daily session (where both round 1 and round 3 were attended) was 91.1% (1241 correct / 1362 total encounters).

The overall combined specificity for all dogs in round 1 of each daily session was 83.6% (766 passed up human control / 916 total human control encounters), and the overall combined specificity for all dogs in round 3 of each daily session was 88.1% (749 passed up human control / 850 total human control encounters).

In both sensitivity and specificity, there was a statistically significant increase of 4.9% (p = 0.0001) and 4.5% (p = 0.007), respectively, between round 1 and round 3 (Table 7). This could indicate that the dogs were becoming more familiar with the PD odor, or that they were becoming more familiar with the procedure of the trials throughout the day.

**Table 7.** Comparison of matched rounds 1 and 3 for all dogs.

| **Sensitivity** | **Number Encounters** | **% Successful** | **95% CI (Percent)** | **Chi-Squared** | **Significance** |
| --- | --- | --- | --- | --- | --- |
| **Round 1** | 1362 | 86.2 | 84.3–88.0 |  |  |
| **Round 3** | 1362 | 91.1 | 89.5–92.6 |  |  |
| **Difference** | | 4.9 | 2.5–7.3 | 16.38 | $p = 0.0001$ |
| **Specificity** |  |  |  |  |  |
| **Round 1** | 916 | 83.6 | 81.1–86.0 |  |  |
| **Round 3** | 850 | 88.1 | 85.8–90.2 |  |  |
| **Difference** | | 4.5 | 1.2–7.7 | 7.30 | $p = 0.007$ |

The location of the PD-positive sample was randomized between each round, so the increase between rounds 1 and 3 was not likely due to memorizing the location of the PD sample. This further denotes that the dogs were not identifying PD-positive samples by memorization.

When comparing sensitivity and specificity for the first round of all unique samples to the final round of the day for all unique samples, the differences were not significant at the 95% level (p = 0.3 and p = 0.96, respectively), as shown in Table 8.

**Table 8.** Comparison of sensitivity and specificity for first and final rounds of the day for all unique samples on the wheel.

| Sensitivity | Number Encounters | % Successful | 95% CI (Percent) | Chi-Squared | Significance |
| --- | --- | --- | --- | --- | --- |
| Round 1 Unique | 161 | 86.3 | 80.1–91.2 |  |  |
| Final Round Unique | 161 | 90.1 | 84.4–94.2 |  |  |
| Difference | | 3.7 | -3.4–10.9 | 1.07 | $p = 0.30$ |
| Specificity |  |  |  |  |  |
| Round 1 Unique | 136 | 89.0 | 82.5–93.7 |  |  |
| Final Round Unique | 111 | 89.2 | 81.9–94.3 |  |  |
| Difference | | 0.2 | -8.08–8.03 | 0.003 | $p = 0.99$ |

Tables 7 and 8 suggest that the dogs were not memorizing target odor by sample or position of sample regardless of whether they were encountering all samples on the wheel for the first time or had encountered them in a prior round.

### Average sensitivity/specificity for the dogs by breed

To show variations in breed groups represented, dogs were categorized as shown in Table 9. These groupings are based on AKC-recognized breed standards (38).

**Table 9.** Combined average percentage sensitivity and specificity for breed groups.

| Breed Group | Number of Dogs | % Sensitivity | Total Rounds | % Specificity | Total Rounds |
| --- | --- | --- | --- | --- | --- |
| Herding | 3 | 91.3 | 495 | 90.9 | 297 |
| Hound | 1 | 62.3 | 61 | 55.3 | 47 |
| Non-Sporting | 4 | 91.9 | 731 | 91.1 | 540 |
| Sporting | 8 | 90.3 | 1525 | 87.5 | 1040 |
| Terrier | 4 | 88.8 | 1269 | 86.4 | 769 |
| Toy | 1 | 81.5 | 298 | 73.3 | 176 |
| Working | 2 | 82.8 | 174 | 81.5 | 124 |

The combined average sensitivity and specificity for the breed groups are reported in Table 9. In most cases, there are not enough dogs in any breed group to make any inferences about particular breed sensitivity or specificity trends. In addition, each dog was raised in a different environment, and most breed groups represented in the study consist of dogs of dissimilar ages with varied days and duration of program attendance.

Figure 3 represents the overall sensitivity for each dog grouped by breed, labeled with dog age. Figure 4 represents the overall specificity for each dog grouped by breed, labeled with dog age. Both Figures 3 and 4 show that breed did not significantly influence outcomes for sensitivity and specificity for this study.

**Figure 3.**
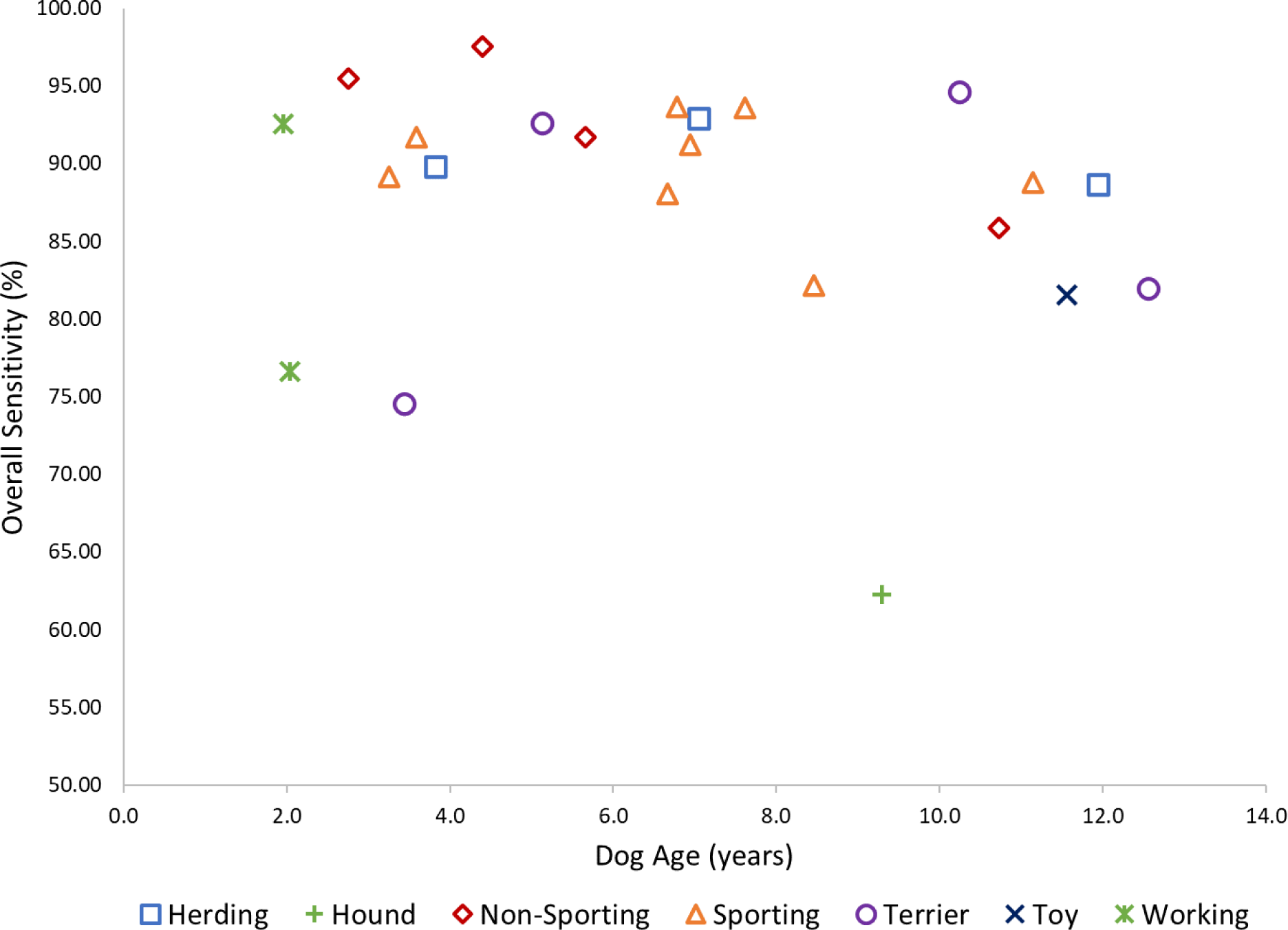
Overall sensitivity for each dog by age and breed.

**Figure 4.**
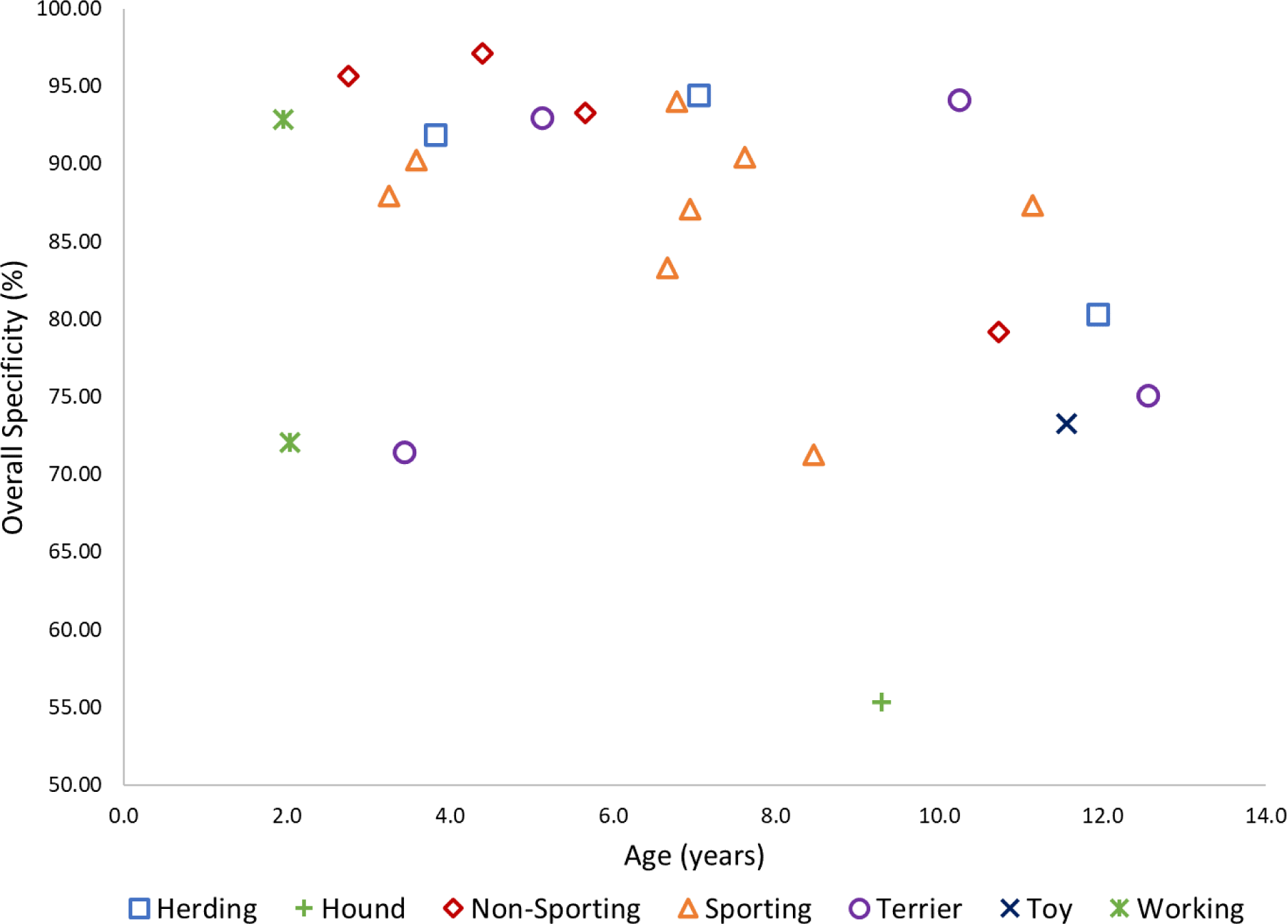
Overall specificity for each dog grouped by age and breed.

### Average sensitivity/specificity compared to total exposures

Figure 5 plots the average sensitivity for each dog during the 2020–2021 study period against the total exposures that each dog had to PD-Positive odor over the life of the PADs program. While there was one dog with low total exposures and low average percent sensitivity, this is likely a special case related to the individual dog. Overall, more total PD-Positive exposures do not produce a higher average sensitivity for any given dog (R^2^ = 0.043, p = 0.344).

**Figure 5.**
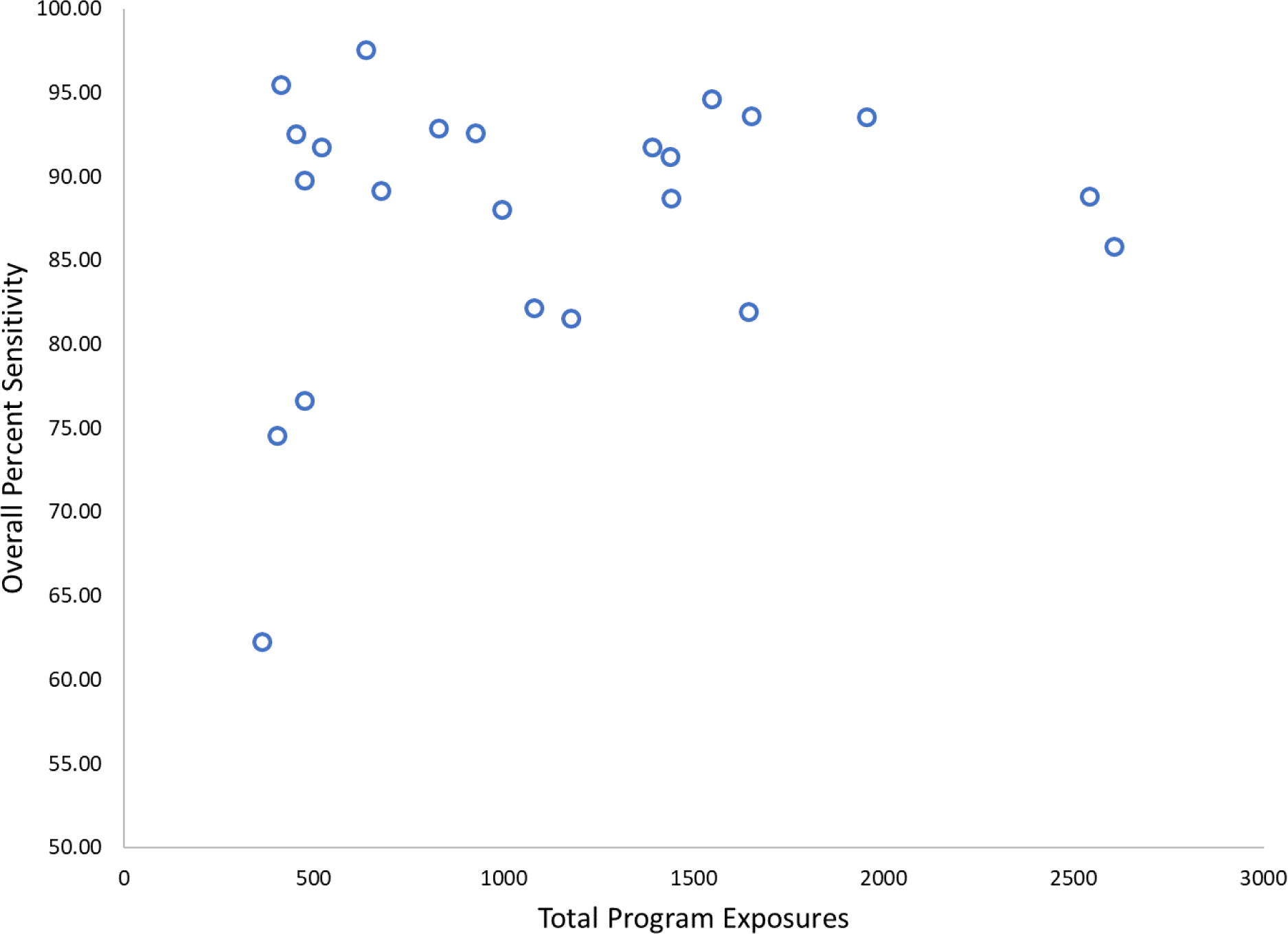
Average percent sensitivity vs. total program exposures for each dog.

Figure 6 plots the average specificity for each dog during the 2020–2021 study period against the total exposures that each dog had to PD-positive odor over the life of the PADs program. Overall, more total PD-positive exposures do not produce a higher average specificity for any given dog (R^2^ = 0.007, p = 0.711).

**Figure 6.**
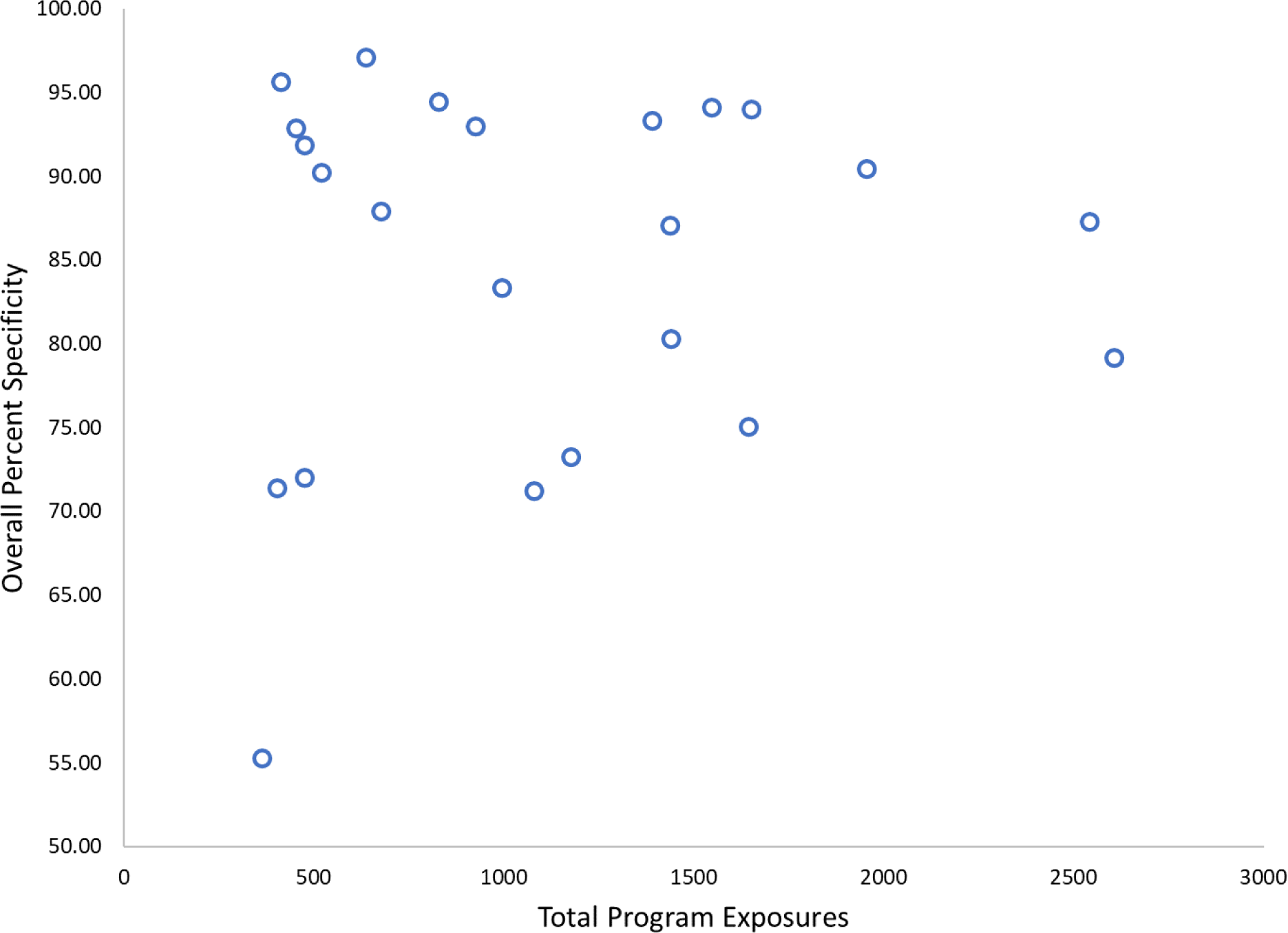
Average percent specificity vs. total program exposures for each dog.

### Sensitivity/specificity of dogs on drug-naïve PD-positive sample participants vs. participating sample donors who report levodopa usage

Overall sensitivity for participating PD-positive sample donors who reported levodopa usage at the time of sample collection was 89.0% (3020 of 3395, 95% CI 87.9% to 90.0%), and sensitivity for participating PD-positive sample donors who reported no levodopa usage at the time of sample collection was 88.25% (954 of 1081, 95% CI 86.18% to 90.11%). Using a two-proportion z-test these percentages are not shown to be significantly different (z = 0.64, two-tailed p-value = 0.52).

Comparing specificity between participating PD-positive sample donors who reported levodopa usage at the time of sample collection vs. those who did not use levodopa showed similar non-significant results. Overall specificity for PD-positive levodopa users was 85.6% (1575 of 1839, 95% CI 84.0% to 87.2%), and specificity for PD-positive non-levodopa users was 87.1% (600 of 689, 95% CI 84.4% to 89.5%). Using a two-proportion z-test, these percentages are not significantly different from each other (z = - 0.69, two-tailed p-value = 0.49). This shows that the dogs were not indicating based on levodopa.

### Comparison of sensitivity/specificity between male PD-positive donors to female PD-positive donors

Table 10 shows the overall sensitivity and specificity for male PD-positive sample donors compared to those of female PD-positive sample donors. For both sensitivity and specificity, there were no significant differences between male PD-positive donors and female PD-positive donors.

**Table 10.** Comparison of overall sensitivity and specificity for male PD-positive sample donors and female PD-positive sample donors.

| **Sensitivity** | **Number Encounters** | **Number Successful** | **% Successful** | **95% CI (Percent)** | **Chi-Squared** | **Significance** |
| --- | --- | --- | --- | --- | --- | --- |
| **Male Donors** | 3553 | 3158 | 88.9 | 87.8–89.9 |  |  |
| **Female Donors** | 1013 | 899 | 88.7 | 86.6–90.6 |  |  |
| **Difference** | | | 0.14 | -2.0–2.5 | 0.015 | $p = 0.90$ |
| **Specificity** |  |  |  |  |  |  |
| **Male Donors** | 2288 | 1974 | 86.3 | 84.8–87.7 |  |  |
| **Female Donors** | 667 | 580 | 87.0 | 84.2–89.4 |  |  |
| **Difference** | | | -0.68 | -2.4–3.5 | 0.20 | $p = 0.65$ |

### Sensitivity/specificity showing the transition from T-shirt samples to swab samples

During September 2022, the samples presented to the dogs transitioned from T-shirts to cotton swabs that had been rubbed over the skin near the back of the neck for both human control and PD-positive donors. Figures 7 and 8 show the daily average sensitivity and specificity trends for the transition period, August 1, 2022, to November 17, 2022.

**Figure 7.**
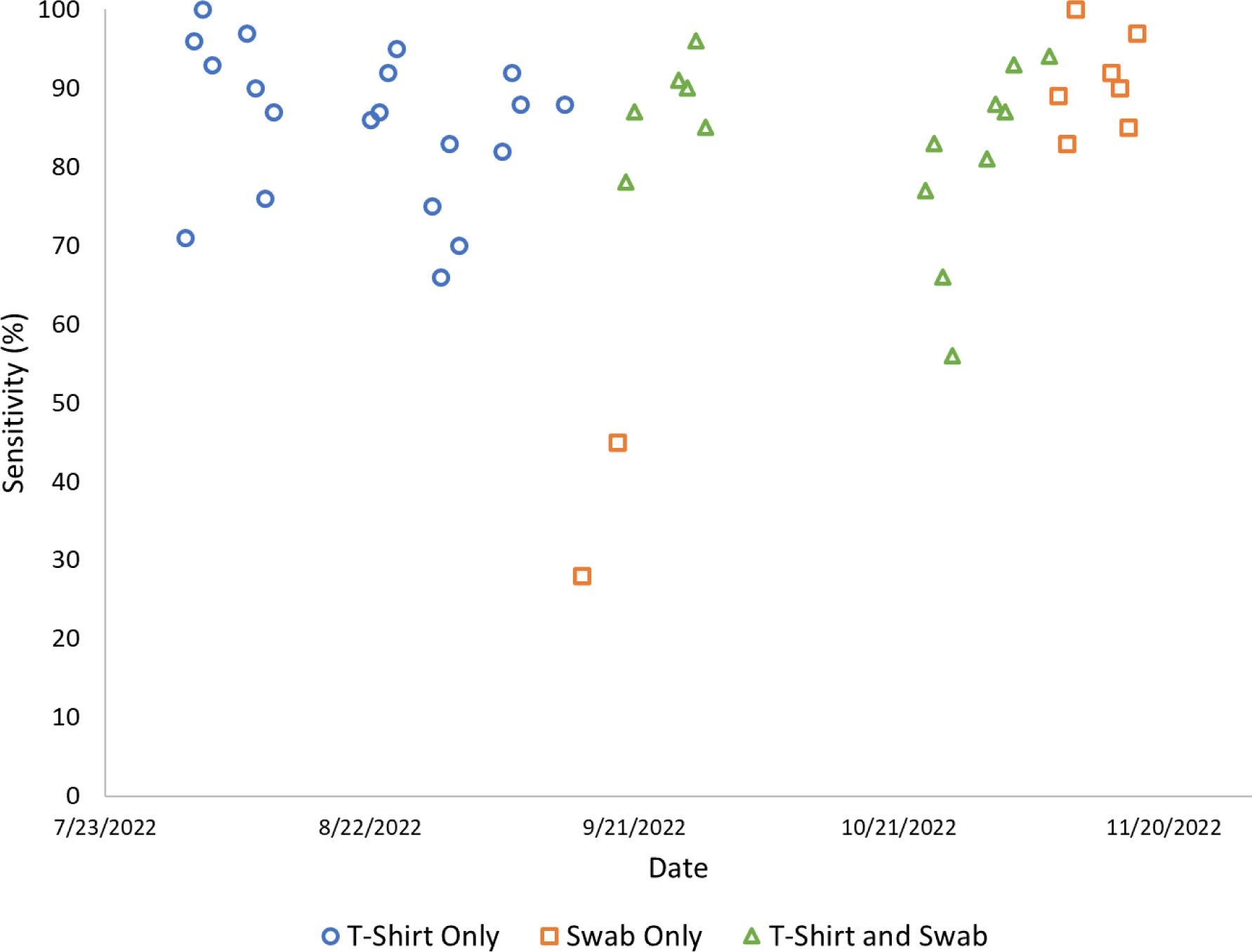
Sensitivity trend during the period of transition from t-shirt samples to swab samples, August 1, 2022, to November 17, 2022. Comparing the seven days prior to the start of the transition from t-shirt samples (blue circles) to swab samples (orange squares) to the seven days after the transition was complete, average percent sensitivity increased from 82.4 (n = 204) to 91.1% (n=180), p = 0.0124.

**Figure 8.**
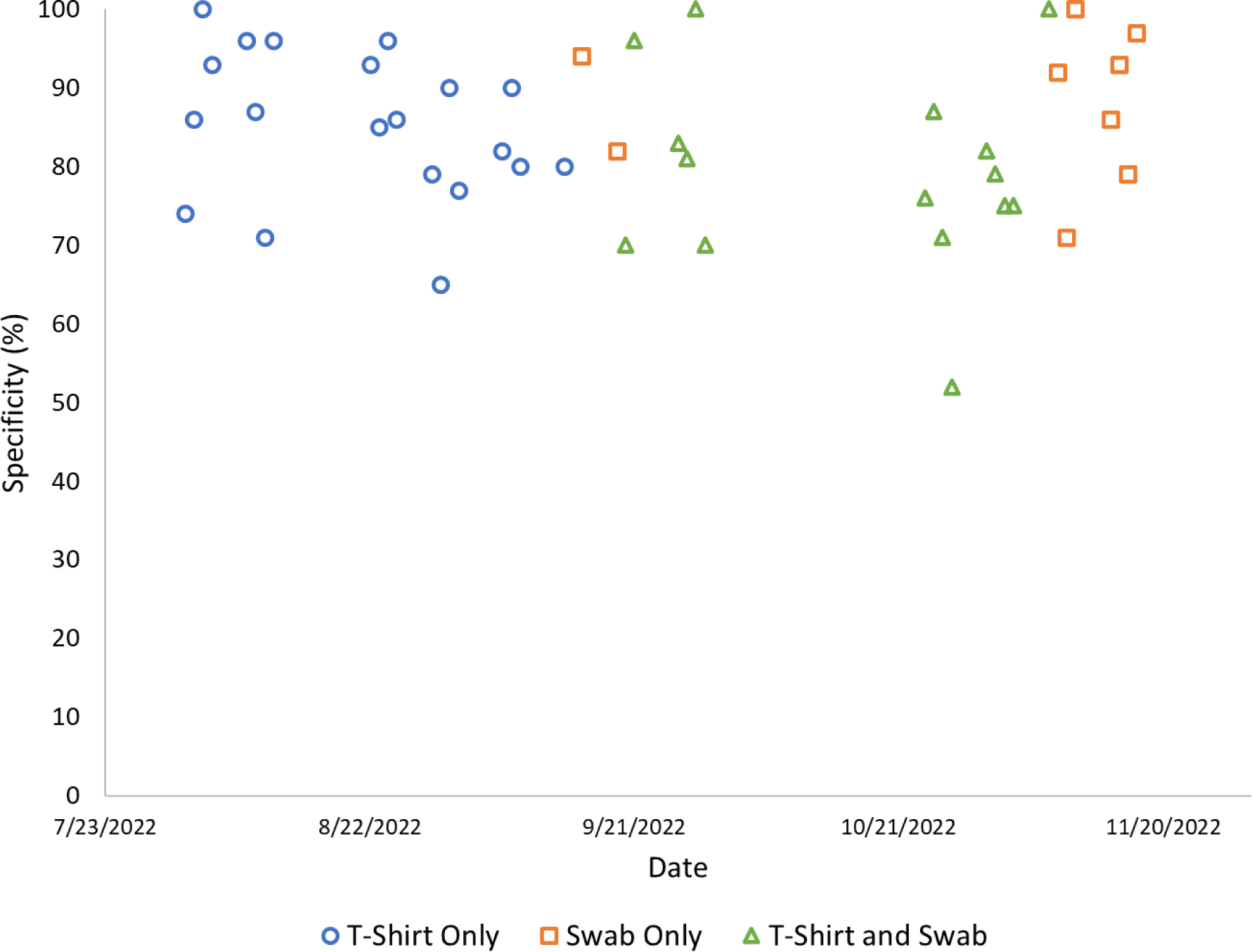
Specificity trends during the period of transition from t-shirt samples to swab samples, August 1, 2022, to November 17, 2022. Comparing the seven days prior to the start of the transition from t-shirt samples (blue circles) to swab samples (orange squares) to the seven days after the transition was complete, average percent sensitivity increased from 81.7 (n = 180) to 88.4% (n=112), p = 0.1256.

During the introduction of cotton swabs only, we saw a significant decline in sensitivity and specificity during September 2022. Once paired with a small piece of T-shirt from the same donor, the results were comparable to the T-shirt-only sensitivity. By November 2022, the dogs showed confidence and proficiency in sensitivity using cotton swabs alone. The decline in sensitivity of September 2022 was included in all reported data. Specificity showed no decline during this period, suggesting that the dogs were not challenged by the absence of PD-associated odor in the sample material. This suggests that for the dogs, the absence of their target odor was understood by them, regardless of the difference in sample material.

## DISCUSSION

In this study, we were able to determine that most household companion dogs when trained through classical detection methods could be used to distinguish between PD-positive and PD-negative samples with a sensitivity of 85% or greater. The top-tier group of dogs was able to distinguish and select PD-positive samples with a sensitivity of 94%. This indicates the selection of top-tier (highly motivated for detection purposes independently of breed) canines would be better suited for the detection of PD-associated odor.

An aspect of canine medical detection training that could be called into question is that of memorization of sample odor so that it could be surmised that target samples that are repeatedly presented to the dogs would realize an increase in sensitivity. In this study, we purposely presented the dogs with samples never encountered and compared these outcomes with samples previously encountered. We repeated this in a high number of instances and under varied conditions. For all the dogs, in all instances, there was less than a 4% difference in sensitivity and specificity between presenting the dogs with “unique” or previously never encountered samples, and samples that had been previously encountered. Findings were consistent even when all the samples presented in the round were unique. This would suggest that the dogs, as a group, were primarily reliant upon their olfactory sensitivity as opposed to any prior familiarization with a specific sample. We further suggest that in determining the sensitivity of canine detection, it would be of value to present findings of the dogs when they encounter all unique donor samples.

An additional aspect of canine detection of PD that is frequently called into question is how drug usage of sample donors affects sensitivity findings. In our study, we found that levodopa usage (the most prescribed drug in PD), did not significantly affect the sensitivity or specificity of the dogs. In Trivedi, et. al, a study that investigated levodopa in sebum secretions, it was determined that changes in sebum from PD-positive patients were not associated with this medication (39). In both Gao (26) and Rooney (27), it was determined that sensitivity and specificity findings were not influenced by drug usage of sample donors.

We did not find that sensitivity and specificity averages for the dogs increased with time spent in training beyond 300 exposures. The study data showed that once the dogs had been in training for one to two years and had achieved between 300 and 600 exposures during that time, the dogs did not increase in their sensitivity and would level off with some slight increases and decreases. Two of the dogs that had been in the program for more than five years showed a marked decrease in sensitivity as they aged, but both dogs had physical disorders—one dog had repeated seizures and the other had a malignant tumor.

Findings over time showed that very few of the dogs were able to recover on the same day from rounds in which reinforcement was not delivered. This was especially true of instances in which the dogs were likely (80% or higher probability) correct in their indication of PD-associated odor and did not receive a reinforcer. These were samples where the sample donor presented with symptoms, but the diagnosis was not by a neurologist, or the diagnosis was more recent than two years. In these instances, the dogs were simply observed in their indication of all unique samples on the wheel in the first round, and not reinforced for their indication unless a total of eight or more of ten dogs had presented with indication on the PD-possible sample.

The decision to not reinforce the dogs for samples in this situation was made based on the science and principles of operant conditioning. Operant conditioning behaviors that are reinforced are more likely to be repeated than behaviors that are not reinforced. For this reason, the dogs, as a group, were then presented with several days of recovery training in which they were presented with samples of known status. This was necessary to rebuild canine confidence that may have been diminished through a lack of reinforcement for a correct response. In the absence of recovery training, the dogs as a group were observed to trend downward in sensitivity. This is likely because companion dogs that are trained with operant conditioning and subjected to increased pressure are more likely to offer behavior leading to false positives in training. In the Hackner study (16), which simulated diagnostic screening conditions for dogs for lung cancer, the specificity of the dogs dropped to 34% under screening conditions, suggesting a high incidence of false positives. The dogs in the study had been trained and were performing at a sensitivity rate of 90% or higher before being subjected to a simulated diagnostic screening test (16,40).

For this reason, we would not recommend the use of household companion dogs for production-style diagnostic screening for PD. Since there is no readily available diagnostic test for the presence of the PD odor, it would not be feasible to reinforce the dogs for every positive indication with confidence in canine accuracy. It requires a high number of dogs to provide statistical significance for sensitivity and, until statistical probability is determined, the dogs cannot be reinforced without risking reinforcement of incorrect behavior. This limits the number of samples the dogs can be used to screen since reinforceable recovery training is needed between the presentation of questionable samples. However, the dogs could be used to effectively pre-select low numbers of samples, working in tandem with complimentary scientific methods to determine if the PD odor is present. This has been useful for the development of electronic or bioelectronic noses (41), for further research into the nature of PD-specific odor, and possibly to understand the evolution of odor as it relates to disease progression.

In Gao et al. (26) it was noted that the dogs in the study from China did not indicate on three separate samples provided by sample providers with a type of PD that is ascribed to a genetic mutation. In our study, it was also noted that none of the dogs indicated on a sample that was provided by a PD-positive donor who reported a GBA gene mutation. We also noted that the dogs in this study did not indicate on a prior confirmed canine-assessed sample from a sample donor who died within two weeks of providing an additional sample to the program. Samples from this donor had been provided to the dogs in prior instances and, in all previous cases, the dogs had indicated, as a group, that the sample was likely PD-positive. For the sample provided two weeks before death, none of the ten dogs in the round indicated a behavioral response to the sample. This would suggest that the target odor for this sample was not present immediately before the death of this sample donor. This could be due to the absence of target odor or qualitative difference in odor volume. Both possibilities require further investigation.

In Gao et al. (26) the researchers used a determination of two of the three dogs providing a sensitive indication to a PD-donor sourced sample to determine the sample as “PD-positive” to assess the sensitivity accuracy of the dogs. In our study, no weighting was done based on the performance of the dogs. In cases where a sample donor had not been diagnosed by a neurologist or had been diagnosed for less than two years, the researchers required the sensitivity of 80% of the dogs in the first daily working session for that sample to receive a determination of PD-positive.

To the best of our knowledge, this is the first detection program and most comprehensive study to investigate the sensitivity of household companion dogs for the detection of PD. This study systematically considered variables that could affect canine detection outcomes including breed, training duration, attendance days, age, and environmental background. Despite these variables, the dogs, as a group, were consistent in their ability to distinguish between PD-positive and PD-negative samples with a sensitivity rate of 89% or higher.

This study serves to advance the field of canine detection of PD as follows:

- Household-raised companion dogs of varying breeds and backgrounds can be used for PD detection, thus eliminating the need for the stewardship responsibility and expense of canine ownership by a detection program.
- Household-raised companion dogs can be used to assess small sample sizes for PD odor detection provided they are given recovery training periods.
- This study provides additional substantiating evidence for the presence of one or more volatile organic compounds obtained from sebum samples of PD-positive patients.
- This study provides additional evidence for the viability of canine detection of PD.

This study demonstrates that companion dogs can detect a Parkinson’s-associated target odor, which likely exists as one or more volatile organic compound(s). If better understood, this odor could be an important biomarker for the early and non-invasive detection of prodromal Parkinson’s disease. If a biomarker can be isolated and reproduced in the form of a singular training aid for dogs, companion dogs could then become a useful, cost-effective, and widespread method of early detection of Parkinson’s disease.

## Competing Interest Statement

The authors have declared that no competing interests exist. This includes any submissions related to patents, patent applications or product development.

## Funding Statement

The author(s) Lisa Holt and Samuel V Johnston received no specific funding for the work of publishing the manuscript.

Lisa Holt was a founding member of the PADs organization, and her salary was paid for through generous contributions of donors.

PADs for Parkinson’s, a 501 (c) (3) nonprofit organization, with the mission of training dogs for the detection of Parkinson’s, funded 5-1/2 years of the seven years of research which included this study. No donors to PADs played a role in the research, authorship, study design, data collection and analysis, decision to publish, or preparation of the manuscript.

## Ethics Statements

This research includes non-human subjects. This research received an IRB exemption (NHS Exemption-Holt (03-14-2023), WCG IRB Work Order #1-1641720-1) under the exclusionary guidelines that the research conducted was for the purpose of canine training for sample selection of PD-associated odor.

## About the Authors

### Lisa Holt

ROLES: PADs for Parkinson’s co-founder, program director, canine detection trainer, study design, writing, original draft, editing, program funding

AFFILIATIONS AND QUALIFICATIONS: Animal Behavior Certified Trainer and Animal Behavior Certified Mentor Trainer, Animal Behavior College; Certified Nose Work Instructor, National Association of Canine Scent Work; Therapy Pet Evaluator, Therapy Pets Unlimited; Certified American Treibball Trainer and Coach, American Treibball Association; Canine Good Citizenship Evaluator, American Kennel Club ORCID ID: https://orcid.org/0000-0002-8212-3375

### Samuel V Johnston

ROLES: Data curation, formal analysis, validation, writing, statistical designer, analyzer, and data presentation

AFFILIATIONS AND QUALIFICATIONS: BS, Fisheries, Cal Poly at Humboldt; Principal Investigator, retired at Innovasea Systems, Inc. ORCID ID: https://orcid.org/0009-0004-1514-4223

## Acknowledgements

ADDITIONAL CONTRIBUTORS: Kim Kyllo-Corson PADs for Parkinson’s research coordinator, sample collection, public relations, data entry, verification, citation references, reviewer; Sandra K Johnston, data entry, verification, citation references, reviewer; Natalie R Johnston, data entry, verification, reviewer; Craig Layman Ph.D., Literature search, editing, publication consultation

PRE-SUBMISSION REVIEWERS: Bryan Bartley Ph.D.; Andy Bary M.S.; Jack P. Bell Ph.D.; Robert P Hawkins Ph.D.; Craig Layman Ph.D.; Peter Ledochowitsch Ph.D.

PADS BOARD MEMBERS: Richard Lind; Sherry Smith Bell; Mark Hopkins; Lori Stokes; Katy Barsamian; Barbara Wright

ADVISORS TO THE BOARD: Jack P. Bell Ph.D.; Carolyn Haugen; Nancy Jones, Co-Founder of PADs

LEGAL COUNSEL: special projects, Markham Quehrn

DOGS AND HANDLER TEAMS: John and Allison Moalli - ‘Bendy’; Leah McConnell - ‘Bubba’; Katy Barsamian - ‘Ella’; Dr. Tim Kopet and Judith Okulitch - ‘Hudson’; Sandy and Natalie Johnston - ‘Indy’; Jack P. Bell Ph.D. - ‘Jaden’; Barbara Wright - ‘Mia and Sasha’; Dick Coffey - ‘Penny’; Bill Moore - ‘Rudi’; Kat Rose - ‘Scarlett’; Amber Chenoweth - ‘Shugga’; Liz Pillow - ‘Suki’; Holly Harbors - ‘Topper’

## Notes

### Competing Interest Statement

The authors have declared no competing interest.

## References

1. Maserejian N, Vinikoor-Imler L, Dilley A. Estimation of the 2020 Global Population of Parkinson’s Disease (PD). In: MDS Virtual Congress 2020. 2020.

2. Willis AW, Roberts E, Beck JC, Fiske B, Ross W, Savica R, et al. Incidence of Parkinson disease in North America. NPJ Parkinsons Dis. 2022;8(1).

3. Rizzo G, Copetti M, Arcuti S, Martino D, Fontana A, Logroscino G. Accuracy of clinical diagnosis of Parkinson disease. Neurology. 2016;86(6).

4. Hess CW, Okun MS. Diagnosing Parkinson Disease. Vol. 22, CONTINUUM Lifelong Learning in Neurology. 2016.

5. Rees RN, Noyce AJ, Schrag A. The prodromes of Parkinson’s disease. European Journal of Neuroscience. 2019;49(3).

6. Kobylecki C. Update on the diagnosis and management of Parkinson’s disease. Clinical Medicine, Journal of the Royal College of Physicians of London. 2020;20(4).

7. Le W, Dong J, Li S, Korczyn AD. Can Biomarkers Help the Early Diagnosis of Parkinson’s Disease? Vol. 33, Neuroscience Bulletin. 2017.

8. https://www.medicaldetectiondogs.org.uk/ [Internet]. Medical Detection Dogs, U.K.

9. Morgan J. Joy of super smeller: Sebum clues for PD diagnostics. Vol. 15, The Lancet Neurology. 2016.

10. Sarkar D, Sinclair E, Lim SH, Walton-Doyle C, Jafri K, Milne J, et al. Paper Spray Ionization Ion Mobility Mass Spectrometry of Sebum Classifies Biomarker Classes for the Diagnosis of Parkinson’s Disease. JACS Au. 2022;2(9).

11. Kokocińska-Kusiak A, Woszczyło M, Zybala M, Maciocha J, Barłowska K, Dzięcioł M. Canine olfaction: Physiology, behavior, and possibilities for practical applications. Animals. 2021 Aug 1;11(8).

12. Jenkins EK, DeChant MT, Perry EB. When the nose doesn’t know: Canine olfactory function associated with health, management, and potential links to microbiota. Vol. 5, Frontiers in Veterinary Science. 2018.

13. Amundsen T, Sundstrom S, Buvik T, Gederaas OA, Haaverstad R. Can dogs smell lung cancer? First study using exhaled breath and urine screening in unselected patients with suspected lung cancer. Acta Oncol (Madr). 2014;53(3).

14. Feil C, Staib F, Berger MR, Stein T, Schmidtmann I, Forster A, et al. Sniffer dogs can identify lung cancer patients from breath and urine samples. BMC Cancer. 2021;21(1).

15. Junqueira H, Quinn TA, Biringer R, Hussein M, Smeriglio C, Barrueto L, et al. Accuracy of canine scent detection of non–small cell lung cancer in blood serum. Journal of the American Osteopathic Association. 2019;119(7).

16. Hackner K, Errhalt P, Mueller MR, Speiser M, Marzluf BA, Schulheim A, et al. Canine scent detection for the diagnosis of lung cancer in a screening-like situation. J Breath Res. 2016;10(4).

17. McCulloch M, Jezierski T, Broffman M, Hubbard A, Turner K, Janecki T. Diagnostic accuracy of canine scent detection in early- and late-stage lung and breast cancers. Integr Cancer Ther. 2006;5(1).

18. Kure S, Iida S, Yamada M, Takei H, Yamashita N, Sato Y, et al. Breast cancer detection from a urine sample by dog sniffing: A preliminary study for the development of a new screening device, and a literature review. Biology (Basel). 2021;10(6).

19. Sonoda H, Kohnoe S, Yamazato T, Satoh Y, Morizono G, Shikata K, et al. Colorectal cancer screening with odour material by canine scent detection. Gut. 2011;60(6).

20. Guest C, Pinder M, Doggett M, Squires C, Affara M, Kandeh B, et al. Trained dogs identify people with malaria parasites by their odour. Vol. 19, The Lancet Infectious Diseases. 2019.

21. Otto CM, Sell TK, Veenema TG, Hosangadi D, Vahey RA, Connell ND, et al. The Promise of Disease Detection Dogs in Pandemic Response: Lessons Learned From COVID-19. Disaster Med Public Health Prep. 2023;17(1).

22. Meller S, Al Khatri MSA, Alhammadi HK, Álvarez G, Alvergnat G, Alves LC, et al. Expert considerations and consensus for using dogs to detect human SARS-CoV-2-infections. Vol. 9, Frontiers in Medicine. 2022.

23. Grandjean D, Gallet C, Lecoq-Julien C, Sarkis R, Muzzin Q, Roger V, et al. SARS-COV-2 Virus Infected Patient Identification Through Canine Olfactive Detection on Axillary Sweat Samples. medRxiv. 2021;

24. Devillier P, Gallet C, Salvator H, Lecoq-Julien C, Naline E, Roisse D, et al. Biomedical detection dogs for the identification of SARS-CoV-2 infections from axillary sweat and breath samples∗∗. J Breath Res. 2022;16(3).

25. Catala A, Grandgeorge M, Schaff JL, Cousillas H, Hausberger M, Cattet J. Dogs demonstrate the existence of an epileptic seizure odour in humans. Sci Rep. 2019;9(1).

26. Gao CQ, Wang SN, Wang MM, Li JJ, Qiao JJ, Huang JJ, et al. Sensitivity of Sniffer Dogs for a Diagnosis of Parkinson’s Disease: A Diagnostic Accuracy Study. Movement Disorders. 2022;

27. Rooney N, Trivedi D. Trained dogs can detect the odour of Parkinson’s Disease. MedRxiv Preprint. 2023;

28. Hall NJ, Glenn K, Smith DW, Wynne CDL. Performance of Pugs, German Shepherds, and Greyhounds (Canis lupus familiaris) on an odor-discrimination task. J Comp Psychol. 2015;129(3).

29. Abbott A. Levodopa: The story so far. Vol. 466, Nature. 2010.

30. Essler JL, Wilson C, Verta AC, Feuer R, Otto CM. Differences in the Search Behavior of Cancer Detection Dogs Trained to Have Either a Sit or Stand-Stare Final Response. Front Vet Sci. 2020;7.

31. Trevethan R. Sensitivity, Specificity, and Predictive Values: Foundations, Pliabilities, and Pitfalls in Research and Practice. Front Public Health. 2017;5.

32. Clopper CJ, Pearson ES. The Use of Confidence or Fiducial Limits Illustrated in the Case of binomial. Biometrika. 1934;26(4):404–13.

33. https://sample-size.net/confidence-interval-proportion/ [Internet]. Confidence interval for a proportion.

34. Campbell I. Chi-squared and Fisher–Irwin tests of two-by-two tables with small sample recommendations. Stat Med. 2007 Aug 30;26(19):3661–75.

35. Richardson JTE. The analysis of 2 × 2 contingency tables—Yet again. Stat Med. 2011 Apr 15;30(8):890–890.

36. MedCalc’s Comparison of proportions calculator [Internet]. [cited 2023 Dec 26]. Available from: https://www.medcalc.org/calc/comparison_of_proportions.php

37. Two Proportion Z-Test Calculator - Statology [Internet]. [cited 2023 Dec 26]. Available from: https://www.statology.org/two-proportion-z-test-calculator/

38. AKC Dog Breed Groups: Understanding Breed Groups at AKC Shows [Internet]. [cited 2023 Dec 26]. Available from: https://www.akc.org/expert-advice/lifestyle/7-akc-dog-breed-groups-explained/

39. Trivedi DK, Sinclair E, Xu Y, Sarkar D, Walton-Doyle C, Liscio C, et al. Discovery of Volatile Biomarkers of Parkinson’s Disease from Sebum. ACS Cent Sci. 2019;5(4).

40. Lazarowski L, Krichbaum S, DeGreeff LE, Simon A, Singletary M, Angle C, et al. Methodological Considerations in Canine Olfactory Detection Research. Vol. 7, Frontiers in Veterinary Science. 2020.

41. Shor E, Herrero-Vidal P, Dewan A, Uguz I, Curto VF, Malliaras GG, et al. Sensitive and robust chemical detection using an olfactory brain-computer interface. Biosens Bioelectron. 2022;195.

